# Loss of CD44 re-educates pancreatic cancer-associated fibroblasts modulating their fibrotic and immunosuppressive functions

**DOI:** 10.64898/2025.12.05.692606

**Authors:** Sven Máté Treffert, Yvonne Madeleine Heneka, Julie Martin, Geoffroy Andrieux, Larissa Launhardt, Eliana Stanganello, Steffen Joachim Sonnentag, Lisa-Marie Mehner, Leonel Munoz-Sagredo, Michelle Christ, Véronique Orian-Rousseau

**Author notes:** These authors contributed equally to the manuscript (Sven Máté Treffert, Yvonne Madeleine Heneka).

## Abstract

Pancreatic tumors are characterized by a prominent stroma that makes up to 90% of the tumor. Due to the significant upregulation of CD44, a family of transmembrane glycoproteins, in pancreatic cancer-associated fibroblasts (CAFs), we investigated its role in myofibroblastic and inflammatory CAF subsets. Conditional deletion of *Cd44* in CAFs in *Cd44^fl/fl^;PdgfrβCreER^T2^* mice, significantly decreased the tumor volume. In human CAFs CRIPSR/Cas9-edited to delete *CD44*, the morphology of the fibroblasts changed drastically: CAFs lost their elongated phenotype and adopted a round shape, reflecting their inactivation. This was accompanied by a significant downregulation of activation markers, unresponsiveness to exogenous stimuli and reduced contractile activity. CD44 absence not only downregulated extracellular matrix proteins in CAFs, thus influencing fibrosis, but also changed the immunomodulatory cytokine secretion. Finally, this inactivation of CAFs upon *CD44* deletion influenced their immunosuppressive effect on dendritic cells (DCs) and on cytotoxic T cells (CTLs), resulting in decreased expression of immunosuppressive cytokines in DCs and enhanced tumor-cell-killing by CTLs.

## Introduction

Pancreatic ductal adenocarcinoma (PDAC), the most common form of pancreatic cancer, arises from the exocrine part of the pancreas [1]. As many patients present with systemic and non-resectable disease, the overall five-year survival rate is low (around 10%) [2]. The aggressive clinical course of PDAC is driven not only by the intrinsic biology of cancer cells, but also by a distinctive tumor microenvironment (TME) that profoundly shapes disease progression and therapeutic resistance [3].

A hallmark of PDAC is the prominent desmoplastic stroma, which can represent up to 90% of the total tumor volume [4] and is composed of extracellular matrix (ECM), immune cells, endothelial cells and cancer-associated fibroblasts (CAFs). Pancreatic stellate cells (PSCs), which are resident mesenchymal cells that store lipid droplets and present fibroblastic features in healthy tissues, have been identified as a possible source of CAFs. Upon exposure to tissue damage or a stiff growth substrate, PSCs become activated and acquire a myofibroblast-like phenotype. Transcriptional analysis of activated PSCs has revealed an upregulation of extracellular matrix components, remodeling enzymes, growth factors, and other signatures characteristic of CAFs [5]. This supports the notion that PSCs are indeed capable of differentiating into CAFs during PDAC progression. Notably, CAFs represent a highly heterogeneous population comprising multiple CAF subpopulations, which arise not only from PSCs but also from bone marrow-derived mesenchymal cells and from epithelial or endothelial cells through transdifferentiation processes [6–9].

Two of the main subclusters observed in previous studies are defined as inflammatory (iCAF) and myofibroblastic (myCAF) CAFs. iCAFs are characterized by low levels of ɑ smooth muscle actin (ɑSMA) and high levels of pro-inflammatory cytokines such as IL-6, CXCL12 and CCL2 [10]. They are mainly induced through the IL-1/Lif/JAK-STAT pathway and the TNF-ɑ pathway, both converging on NF-κB downstream [11, 12]. myCAFs exhibit high ɑSMA expression and produce abundant ECM with an anisotropic arrangement of a variety of extracellular matrix proteins like collagen I, thus acting as a physical barrier that impairs immune cell infiltration and drug delivery in the tumor. Their most potent inducer is transforming growth factor β 1 (TGFβ1) [11].

In addition to their structural role, CAFs exert profound immunomodulatory effects. By secreting a variety of cytokines, chemokines and growth factors, CAFs have been shown in pancreatic and other types of cancer to affect immunosuppressive cells like tumor-associated macrophages (TAMs) [13], myeloid-derived suppressor cells (MDSCs) [14, 15], dendritic cells (DCs) [16, 17] and ultimately T lymphocytes. Several studies have shown that CAFs can affect the identity and function of T cells, thus hindering CD8^+^ cytotoxic T cell (CTL)-mediated responses against the tumor as well as enhancing generation of regulatory T cells (T_reg_) [18–20]. This high abundance of immunosuppressive cell types (MDSCs, TAMs, regulatory T cells), as well as the dampened activity of effector immune cells (CD4^+^ and CD8^+^ T cells), renders pancreatic cancer extremely non-immunogenic and non-responsive to immunotherapy [21].

Given their abundance, plasticity and impact on disease progression, CAFs represent a compelling therapeutic target. However, depleting studies of CAFs in mouse models of pancreatic cancer generated contradictory and controversial results. Many studies found that the depletion of CAFs worsened the outcome, resulting in poorly differentiated and more aggressive tumors [22–24]. Others showed that, depending on the affected population, targeting various subtypes in the distinct subpopulations resulted in tumor-promoting or tumor-restricting outcomes [25]. Therefore CAF-targeting strategies have been approached with increasing caution and the focus shifted toward the reprogramming of CAFs into a lesser tumor-promoting state [26].

Our group previously showed that blocking a protein isoform of the glycoprotein CD44, CD44v6 in pancreatic orthoptopic xenograft and the autochthonous *Kras^G12D/+^;Trp53^R172H/+^;Pdx1Cre* (KPC) mouse models led to a reduction of primary tumor volume and metastatic burden, even if the species-specific blocking peptides only targeted the stromal CD44v6 [27, 28]. This suggested that CD44 expression in stromal cells critically influences the activity of CAFs during pancreatic tumor progression, as they are central regulators of tissue homeostasis and integrity. However, the mechanistic contribution of CD44 to CAF activation, ECM organization, and immunoregulatory capacity in PDAC has remained unclear. Here, we show that CD44 is essential for CAF plasticity, ECM organization, and immune evasion in PDAC and support its potential as a target for stroma-reprogramming strategies aimed at improving anti-tumor immunity.

## Results

### CAF-specific knockout of *Cd44 in vivo* significantly associated with reduced primary tumor volume

Previous studies by our group highlighted the role of stromal CD44 during pancreatic tumor progression [27, 28].To identify which stromal compartment contributes to this effect, we analyzed publicly available single-cell RNA sequencing (scRNAseq) data from pancreatic cancer patients [29]. Comparative analysis of non-tumorous (N1 to N11) and tumorous pancreatic tissue (T1 to T24) (Figure 1A) revealed a significant upregulation of CD44 expression in fibroblastic cells in the tumor samples. This selective upregulation shows a prominent association of CD44 with cancer-associated fibroblasts, suggesting a functional contribution of this cell-surface molecule in the stroma of PDAC.

**Figure 1.**
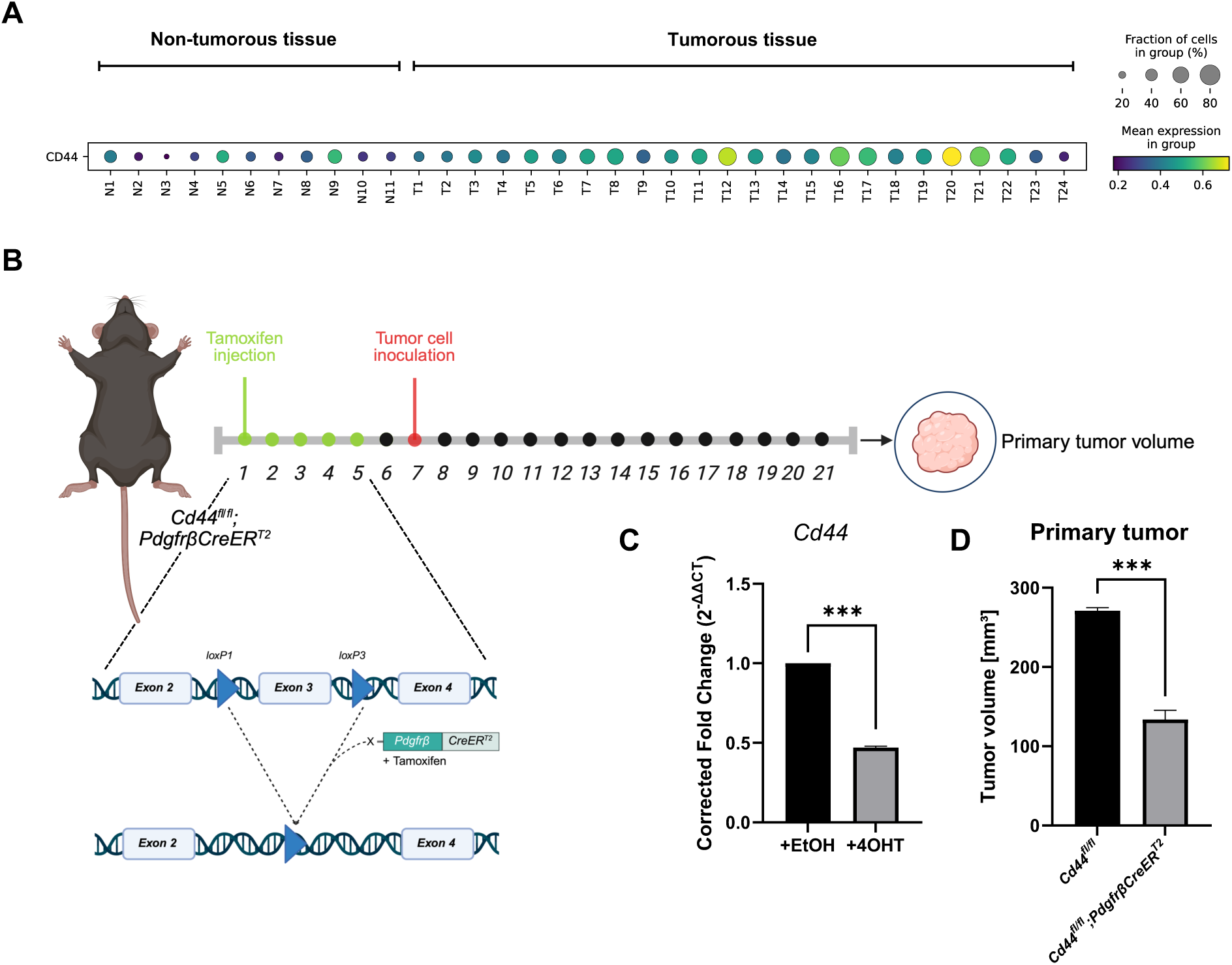
CAF-specific knockout of *Cd44 in vivo* reduces primary tumor volume. **A** Publicly available RNAseq data from human patients was used to compare the expression of CD44 in fibroblastic cells between non-tumorous (N1 to N11) and tumorous pancreatic tissue (T1 to T24). Cell type annotations from the original publication were used as the reference labels. Marker gene analysis was conducted using the Wilcoxon rank-sum test, comparing each cell type against all others. Comparisons between pancreatic cancer and normal conditions were performed using the same approach. *p-value <0.05. **B** *Cd44^fl/fl^;PdgfrβCreER^T2^*mice and the corresponding control animals were injected intraperitoneally with tamoxifen for five days. Two weeks after orthotopic injection of FC1245 cancer cells at day seven, mice were sacrificed and the primary tumor was isolated. **C** Primary pancreatic fibroblasts were isolated from *Cd44^fl/fl^;PdgfrβCreER^T2^*mice and treated with either 25 µM 4-OHT or ethanol as control. *Cd44* expression was analyzed at day seven by qPCR. *Gapdh* and *βactin* were used as reference genes. Data are means ± S.E. Statistical significance was determined using the one sample t-test. n=3. ***p-value<0.005. **D** Primary tumor volume of *Cd44^fl/fl^;PdgfrβCreER^T2^*and corresponding control animals. Data are means ± S.E. Statistical significance was determined using the unpaired one-sided t-test. N=3. ***p-value<0.0002. Parts of the Figure: Created in BioRender. Treffert, S. (2026) https://BioRender.com/efu5df8.

To investigate the potential role of CD44 expression *in vivo*, we used *Cd44^fl/fl^* mice crossed with mice expressing an inducible Cre recombinase (CreER^T2^) under the control of the *Pdgfrβ* promoter (Figure 1B) [30, 31]. This allowed the specific knockout of *Cd44* in cells with an active *Pdgfrβ* promoter, including PSCs and CAFs, through the administration of tamoxifen, which is recognized by the mutated estrogen receptor (ER^T2^). Isolated fibroblastic cells from the pancreas of *Cd44^fl/fl^;PdgfrβCreER^T2^* mice treated with 4-Hydroxytamoxifen (4-OHT) expressed significantly less *Cd44* compared to ethanol-treated control cells (Figure 1C). When they were between 6 weeks and 10 weeks old, male mice of this strain were injected intraperitoneally with tamoxifen (TAM) for 5 days followed by the orthotopic injection of FC1245 cells from the KPC (*Kras^G12D^;Trp53^R172H^;Pdx1-Cre*) mouse model. Strikingly, deletion of *Cd44* in PDGFRβ^+^ stromal cells led to an approximately 50% reduction in primary tumor volume at two weeks after cancer cell implantation compared to control mice (Figure 1D).

### CD44 expression on PSCs supports their TGFβ1-mediated activation into a CAF-like state

The cell populations that were targeted in the *Cd44^fl/fl^;PdgfrβCreER^T2^*mice include CAFs and pancreatic stellate cells (PSCs), known to be activated into a myofibroblastic CAF-like state in response to tumor-derived factors such as TGF-β [5, 11]. To study the influence of CD44 on PSC activation toward a CAFs like state, we used CRISPR/Cas9 to delete *Cd44* in an immortalized murine pancreatic stellate cell line (imPSC) [32] (Figure S1A). We designated these cells as imPSC*^ΔCD44^* cells. We then tested whether the activation properties using TGFβ1 induction and checking the expression of myofibroblastic activation markers COL1A1 and ɑSMA (*Acta2*) (Figure 2A-C). We show that in striking contrast to imPSCs, imPSC*^ΔCD44^* failed to further upregulate these markers in response to TGFβ1. Immunofluorescence analysis revealed that COL1A1 in the imPSCs more than doubled upon induction with TGFβ1, while the increase was significantly less in the imPSC*^ΔCD44^* (Figure 2A). Our qPCR analysis 72 hours after TGFβ1 treatment showed a decrease of *Col1a1* gene expression in the imPSC*^ΔCD44^*, while *Acta2* increased significantly less in comparison to imPSCs (Figure 2B). Furthermore, by western blot we show that TGFβ1 increased the protein level of COL1A1 from 1.0 to 2.03 in the imPSCs, whereas the level of COL1A1 in the imPSC*^ΔCD44^* remained stable (Figure 2C). To exclude CRISPR/Cas9 off-target effects, *Col1a1* expression was analyzed in isolated quiescent pancreatic stellate cells from a *Cd44^fl/fl^;GfapCre* mouse model (Figure S1A). In these mice, *Cd44* is exclusively knocked out in the GFAP-positive quiescent PSCs (Figure S1B). This analysis also revealed a lower expression of *Col1a1* in knockout cells compared control animals upon TGFβ1 treatment (Figure S1C). Consistent results were obtained when CD44 was knocked down via siRNA in the imPSCs (Figure S2A and B). Together, these findings demonstrate that CD44 is required for the TGFβ1-mediated activation of PSCs.

**Figure 2.**
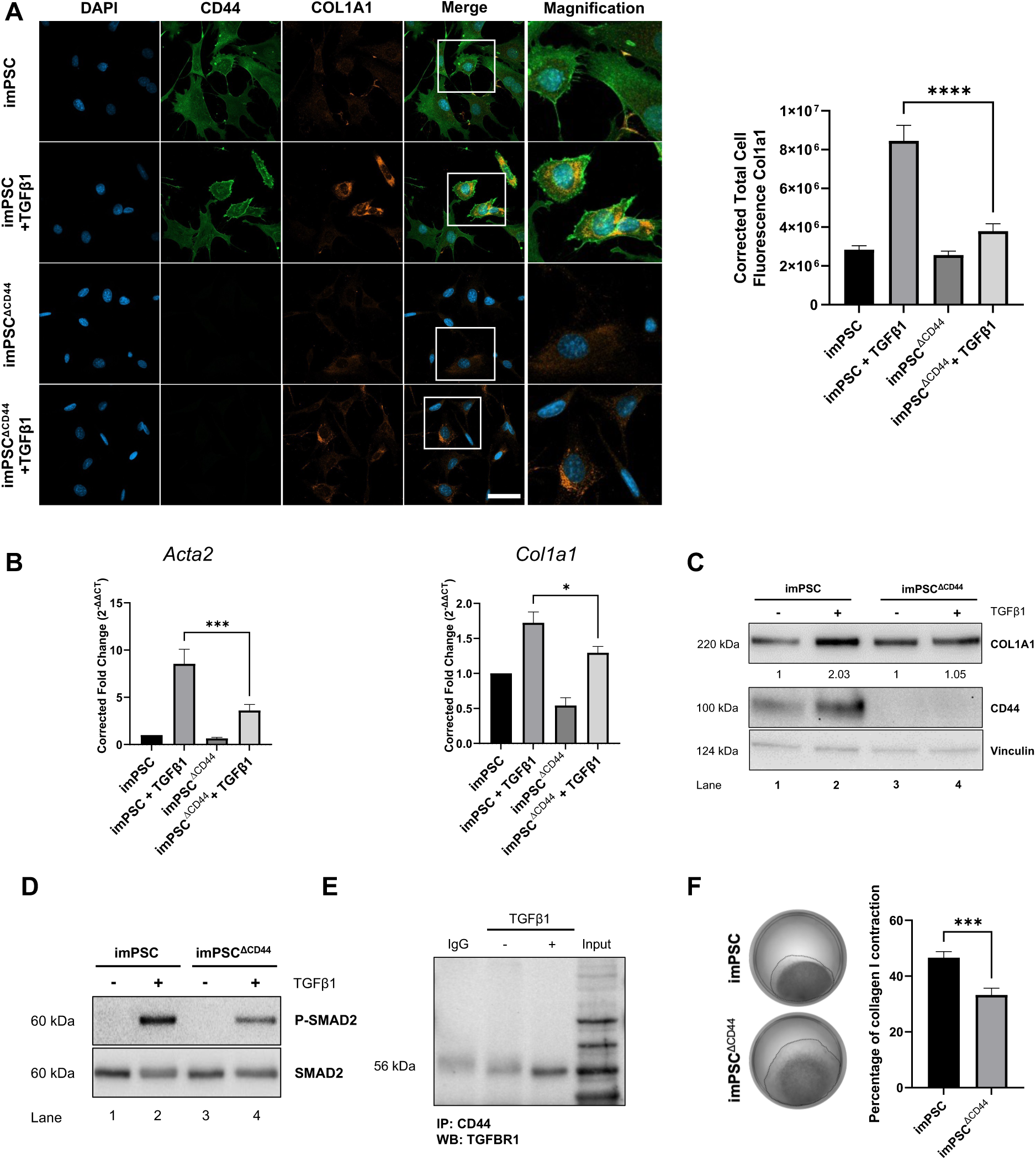
Knockout of *Cd44* in pancreatic stellate cells impairs the induction of a CAF-like state. **A** IF analysis of imPSCs and imPSC*^ΔCD44^* after induction with recombinant TGFβ1 (50 ng/ml) stained for CD44 (AlexaFluor488), ɑ-1 type I collagen (COL1A1, AlexaFluor546); nuclei were counterstained with DAPI. Scale bar, 50 µm. Corrected total cell fluorescence was calculated with ImageJ. N=3, n=30 pictures per condition. Data are means ± S.E. Statistical significance was determined using the one-sided unpaired Student’s t test. **** p-value<0.0001. **B** Gene expression of *Acta2* and *Col1a1* of TGFβ1-induced imPSCs and imPSC*^ΔCD44^* was analyzed by qPCR. *Gapdh* and *βactin* were used as reference genes. N=3 Data are means ± S.E. One-way ANOVA and Holm-Šídák’s multiple comparisons post-hoc test were used for statistical analysis. *p-value<0.0332; ***p-value<0.0021. **C** Analysis of CD44 and COL1A1 on protein level after treatment of imPSCs and imPSC*^ΔCD44^*with TGFβ1 by western blot. Vinculin served as loading control. **D** imPSCs and imPSC*^ΔCD44^* treated 30 minutes with recombinant TGFβ1 (50 ng/ml) were subjected to western blot analysis for P-SMAD2 and SMAD2. **E** Co-immunoprecipitation (Co-IP) of CD44 and TGFβRI in imPSCs. Whole cell lysates were incubated with a CD44 antibody (clone: KM201) or IgG control (rat IgG1,κ) and precipitated. CD44 and TGFβRI were detected via western blot. Input represents non-precipitated lysate. **F** imPSCs and imPSC*^ΔCD44^* were seeded into a 3D type I collagen matrix and cultured for 48 hours. Percentage of contraction was calculated by measuring the area of the contracted gel. N=3. Data are means ± S.E. Statistical significance was determined using the one-sided unpaired Student’s t test. ***p-value<0.00021.

As CD44 proteins act as co-receptors for a variety of cell surface receptors, we assessed whether the knockout of *Cd44* in imPSCs directly influences the TGFβ1-induced Smad pathway, where activation of the receptor leads to phosphorylation of downstream R-SMADs like SMAD2 [33]. We showed that the knockout of *Cd44* led to a reduced phosphorylation of SMAD2 (P-SMAD2) in imPSC*^ΔCD44^* (Figure 2D), indicating that CD44 has a role in TGF-βRI signaling in PSCs. Additionally, co-immunoprecipitation analysis revealed an association between TGF-βRI and CD44 in imPSCs (Figure 2E). The TGFβ1-mediated activation of PSCs also involves EMT-like processes, where mesenchymal markers are upregulated while epithelial markers are decreased [34]. qPCR analysis of TGFβ1-induced imPSCs and imPSC*^ΔCD44^*showed that the knockout of *Cd44* influences these dynamics. This is reflected by a reduced expression of the EMT transcription factors *Snai1*, *Twist1* and *Zeb1* in the imPSC*^ΔCD44^* upon induction by TGFβ1 and a significantly higher expression of *EpCam* (Figure S2C). Using vimentin-chromobody transfected cells, we also confirmed reduced expression of this mesenchymal cytoskeletal protein (Figure S2D) [35].

Based on the findings that the activation into a CAF-like state of PSCs is disturbed by the knockout of *Cd44*, we compared the ability of imPSCs and imPSC*^ΔCD44^* to remodel and contract three-dimensional type I collagen matrices. This is a commonly cited hallmark of *in vitro* fibroblast activation into a myCAF-like state [36, 37] (Figure 2F). After 48 hours of culture in collagen, imPSC*^ΔCD44^*exhibited reduced contractile activity compared to CD44-expressing imPSCs. In summary, we demonstrate that CD44 is required for the TGFβ1-mediated activation of PSCs and contributes functionally to their transition toward a CAF-like state.

### CD44 in pancreatic fibroblastic cells induces tumor stroma-like parallel orientation of collagen I fibers

Activated PSCs and CAFs play a central role in the production, organization and remodeling of the extracellular matrix across various tissue microenvironments, including PDAC. To test the capacity of imPSCs that express or do not express C*d44* to control ECM dynamics, we compared imPSC-/ imPSC*^ΔCD44^*-derived matrices (DM) [38] (Figure 3A). These matrices produced by fibroblastic cells recapitulate key biochemical and architectural features of the *in vivo* TME [39]. After the nine-day generation of a PSC-derived matrix, clear morphological differences became apparent between imPSC*^ΔCD44^* and imPSC control cells following induction into a CAF-like state by TGFβ1 treatment. imPSCs expressing *Cd44* adopted an elongated spindle-like morphology (Figure 3B), and organized into parallel patterns (Figure 3C), as visualized by phalloidin staining. Since tumor-associated ECM synthesized by CAFs typically displays a parallel organization [40], we then investigated whether CD44 expression influences the topography of imPSC-DM by assessing collagen alignment using second harmonic generation (SHG) analysis (Figure 3D). We observed a significant reduction of organized collagen fibers in imPSC*^ΔCD44^*-DM compared to the DM of *Cd44*-expressing imPSCs, reflected in a significant reduction of the SHG intensity. In addition, the level of organization of the assorted imPSC-produced collagen fibers was quantified by measuring the relative orientation angles of fibers [41]. The average percentage of parallel fibers that were oriented within 15° of the mode angle was determined by using the OrientationJ plugin of Fiji. The measured percentages after TGFβ1 induction were 49% for the imPSC and 32% for the imPSC*^ΔCD44^*-DM (Figure 3E). With these results we show that CD44 is necessary for the topographical organization of the ECM fibers produced by imPSCs.

**Figure 3.**
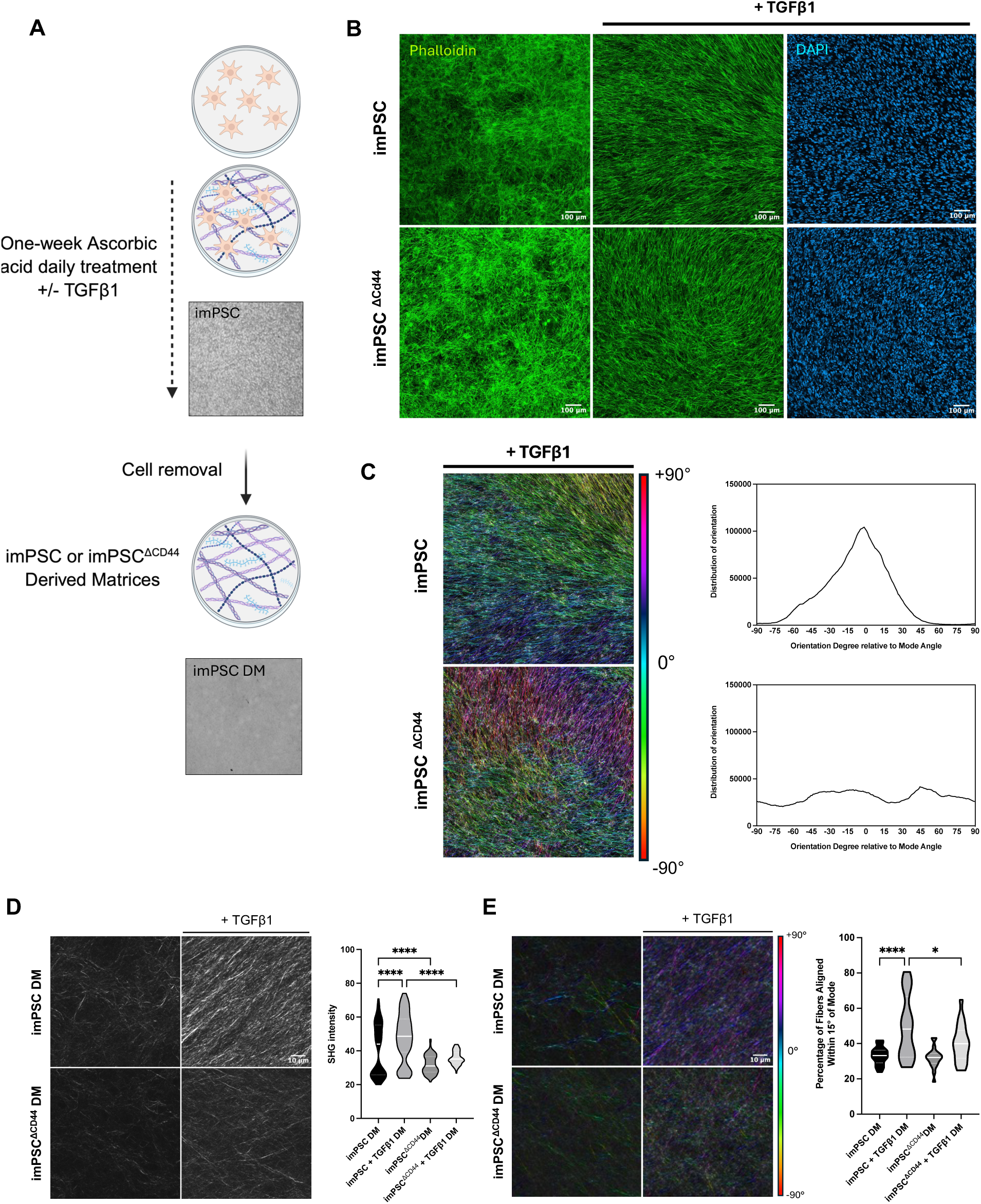
*Cd44* knockout in imPSCs impairs extracellular matrix production and organization. **A** Experimental strategy to generate matrices derived from imPSCs and imPSC*^ΔCD44^*. Schematic illustration: Created in BioRender. Martin, J. (2026) https://BioRender.com/vbggzb6. **B** Representative images of imPSCs and imPSC*^ΔCD44^* during ECM production with or without TGFβ1 stimulation prior to cell removal. Nuclei are stained with DAPI and Phalloidin in green. **C** F-actin fibers digitally pseudo-colored to depict cell orientation angles. The colored bar on the right indicates angle distributions normalized to mode angle (0°, cyan). Cell alignment was quantified by measuring fiber angle distribution. **D** Reconstituted confocal images of Second Harmonic Generation (SHG) microscopy of imPSC- and imPSC*^ΔCD44^*-derived matrices (DM). Quantification of SHG signal intensity reflects the amount of aligned collagen fibers. **E** Analysis of ECM fiber orientation from SHG images in D. Fiber alignment was quantified as the percentage of fibers oriented within 15° of the mode angle. Statistical significance was determined by one-way ANOVA and Holm-Šídák’s multiple comparisons post-hoc test. N=3, n=5 pictures per sample. *p-value<0.0332; ****p-value<0.0001.

### Removal of CD44 from human pancreatic CAFs drastically impacts their activation as reflected by a drastic change in morphology

Our findings above show that *Cd44* deletion in PDGFRβ^+^ cells reduced PSC activation, limited their transition into a myCAF-like state and changed their capacity to produce and remodel the ECM. This attenuation of stromal activity was accompanied by reduced primary tumor growth. To find out whether our findings were translatable to the human context, we examined CD44 function in human tumor-derived CAFs. Using the CRISPR/Cas9 system, we knocked out *CD44* in CAFs from a human pancreatic tumor. Surprisingly, CAFs devoid of *CD44* expression (Figure 4A and Figure S3A), designated as CAF*^ΔCD44^,* lost their typical elongated spindle-like shape and exhibited a small and round morphology. This was not observed in scramble sgRNA-mCherry-transduced control cells (Figure S3B). Because rounded fibroblasts are typically less active [42], we examined their ability to contract three-dimensional type I collagen matrices, analogous to those we tested in the imPSCs. Interestingly, 48 hours after embedment of cells in a collagen matrix, CAF*^ΔCD44^* contracted the gel significantly less compared to *CD44*-expressing controls (Figure 4B), which was similar to our first results on imPSC*^ΔCD44^* (Figure 2F). Both western blot (Figure 4D) and immunofluorescence analyses (Figure S3C) showed a loss or a reduction of key myCAF activation markers COL1A1 and ɑSMA in CAF*^ΔCD44^*, which supports the hypothesis that knocking out *CD44* leads to CAF inactivation. Moreover, we observed that the ɑSMA-containing stress fibers, which are present in CD44-expressing CAFs, are absent in CAF*^ΔCD44^*(Figure S3C).

**Figure 4.**
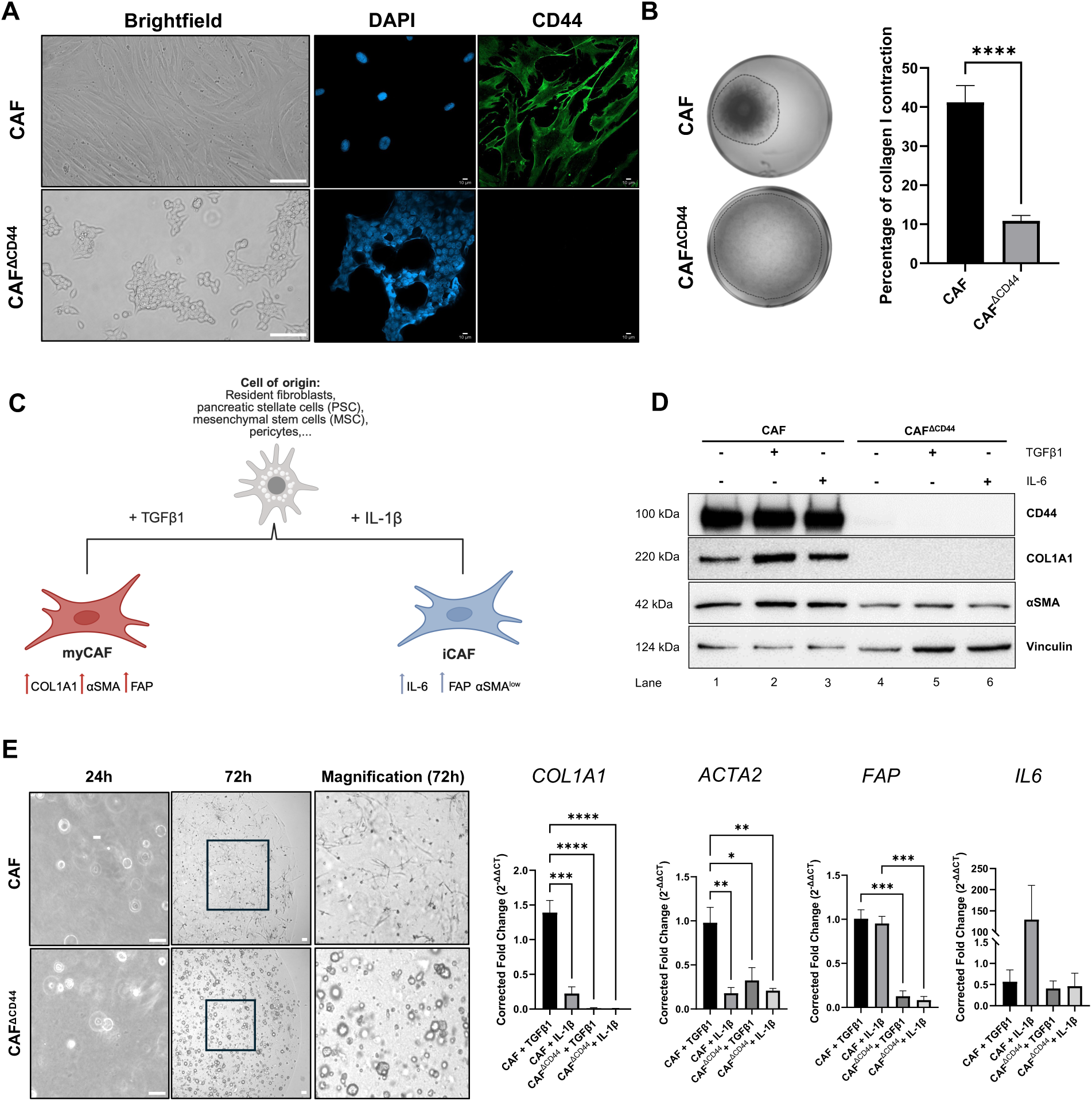
*CD44* knockout in human CAFs influences their morphology, activity and expression of CAF activation markers. **A** Representative image of the morphology of CAFs in comparison to CAF*^ΔCD44^*. Scale bar, 100 µm (left); 10 µm (right). IF staining with AlexaFluor488 visualizes CD44 in CAFs and CAF*^ΔCD44^*. Nuclei were counterstained with DAPI. **B** CAFs and CAF*^ΔCD44^* were seeded into 3D collagen I matrices, incubated for 48 hours, and the percentage of contraction was calculated. N=3. Data are means ± S.E. Statistical significance was determined using the unpaired one-sided Student’s t-test ****p-value<0.0001. **C** The two predominant CAF subtypes, myCAFs and iCAFs, are induced by either TGFβ1 or IL-1β and up- or downregulate their respective activation markers. Schematic illustration: Created in BioRender. Treffert, S. (2026) https://BioRender.com/7t44o62. **D** CAFs and CAF*^ΔCD44^* were treated with recombinant TGFβ1 (50 ng/ml) or IL-6 (20 ng/ml) for 24 hours. Protein expression of CD44, COL1A1 and ɑSMA with Vinculin as loading control. **E** CAFs and CAF*^ΔCD44^* were seeded in Matrigel domes and induced with recombinant TGFβ1 (50 ng/ml) or IL-1β (20 ng/ml) for 24 hours. Representative images of the CAFs and CAF*^ΔCD44^* morphology 24 hours and 72 hours after seeding. Scale bars, 50 µm. *COL1A1*, *ACTA2*, *FAP* and *IL6* gene expression was analyzed by qPCR 24 hours after induction (72 hours after seeding). *GAPDH* was used as a reference gene. N=3. Data are means ± S.E. One-way ANOVA and Holm-Šídák’s multiple comparisons post-hoc test were used for statistical analysis. N=3. *p-value<0.0332; **p-value<0.0021; ***p-value<0.00021; ****p-value<0.0001.

After administering TGFβ1 to the CD44-expressing CAFs, we observed higher COL1A1 protein levels (Figure 4D). However, as seen in the same blot, exogenous TGFβ1 did not lead to the reacquiring of COL1A1 protein in the CAF*^ΔCD44^*, indicating that the knockout of *CD44* in CAFs disturbs the response to TGFβ1.

To investigate the influence of CD44 on the inflammatory CAF subtype (iCAF), CD44-expressing CAFs and CAF*^ΔCD44^* were seeded in Matrigel (Figure 4E), since culturing on stiff surfaces tends to favor the induction and maintenance of the myofibroblastic phenotype [43, 44]. This enabled us to work with cells that were less biased towards a certain subtype. Downregulation of *COL1A1* and upregulation of *IL6* in response to IL-1β confirmed a polarization of CD44-expressing CAFs into iCAFs. CAF*^ΔCD44^* did not upregulate *IL6* and remained at basal RNA level, indicating a disturbed response to IL-1β (Figure 4E). Secretome analysis confirmed a low basal level of IL-6 release by CAF*^ΔCD44^* (Figure 5A). TGFβ1-induced CAF*^ΔCD44^* again presented no *COL1A1* expression along with reduced *ACTA2* (ɑSMA), while both IL-1β and TGFβ1-induced CAF*^ΔCD44^*showed low levels of *FAP*. Interestingly, as observed on culture plastic, CAF*^ΔCD44^* also remained round in Matrigel, while the CD44-expressing CAFs elongated even in Matrigel (Figure 4E). The findings above indicate that the knockout of *CD44* in CAFs isolated from a human tumor reverts their activation and renders them unresponsive to TGFβ1 and IL-1β.

**Figure 5.**
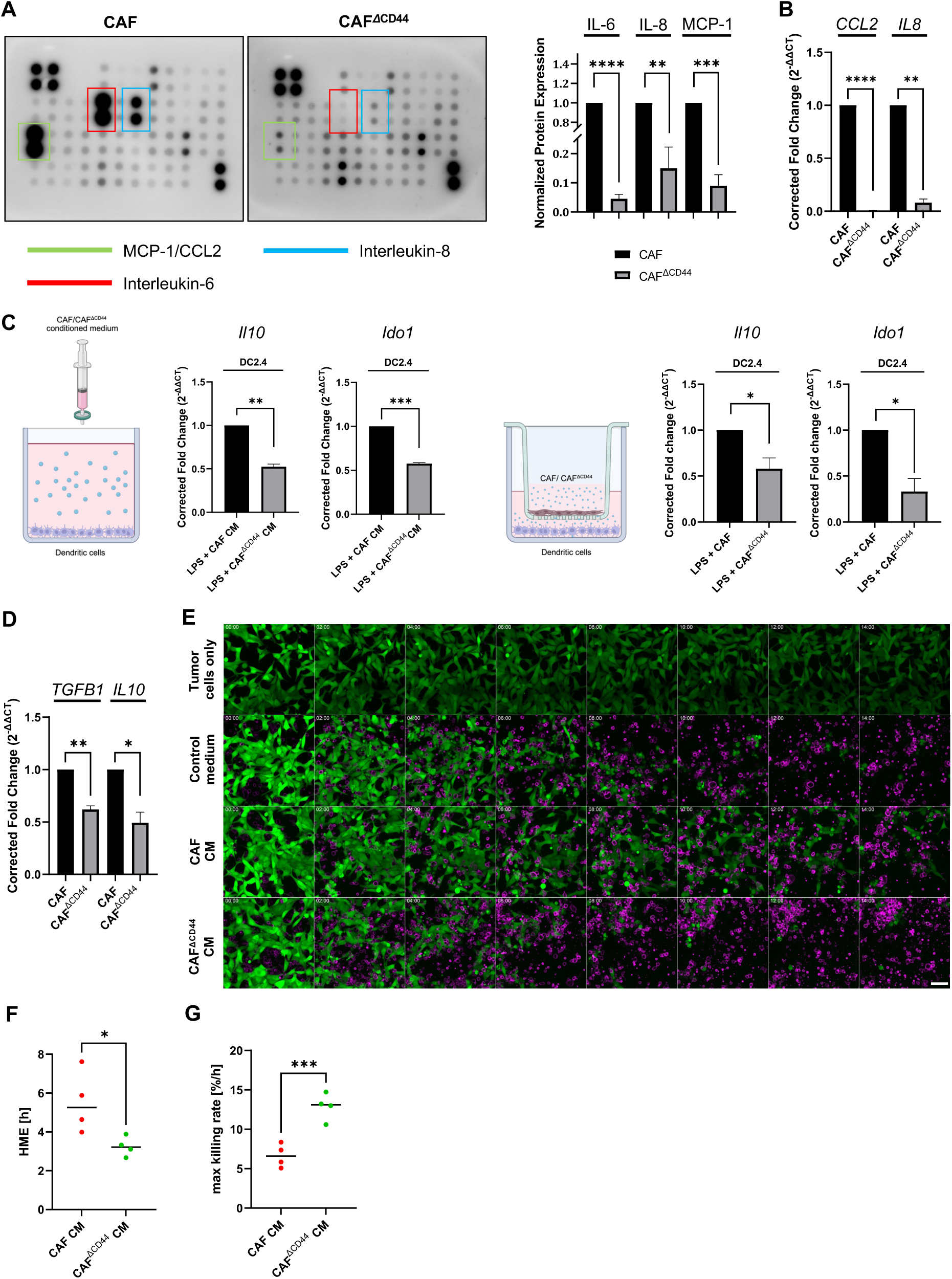
CAF*^ΔCD44^*change their secretome and show a diminished immunosuppressive effect. **A** Conditioned media (CM) from CAFs and CAF*^ΔCD44^* were collected and analyzed *via* cytokine array. Relevant proteins were highlighted by colored rectangles and subsequently quantified using ImageLab software. The mean intensity of the array spots was evaluated and normalized. The normalized protein expression of IL-6, IL-8 and MCP-1 is depicted as ratios comparing CAFs to CAF*^ΔCD44^*. N=3. Data are means ± S.E. Statistical significance was determined using the one sample t-test. **p-value<0.0021; ***p-value<0.0002; ****p-value<0.0001. **B** RNA of CAFs and CAF*^ΔCD44^* was extracted 48 hours after seeding and subjected to qPCR analysis of expression of *CCL2* and *IL8*. *GAPDH* was used as a reference gene. N=3. Data are means ± S.E. Statistical significance was determined using the one sample t-test. **p-value<0.0021; ****p-value<0.0001. **C** DC2.4 were either cultured with 50% CAFs/CAF*^ΔCD44^* conditioned media (CM) or indirectly co-cultured with CAFs/CAF*^ΔCD44^* (as indicated in the illustrations: Created in BioRender. Treffert, S. (2026) https://BioRender.com/ifl9gho) for 48 hours and treated with LPS. At 24 hours after seeding, 1 µg/ml lipopolysaccharide (LPS) was added to the cultures. *Il10* and *Ido1* expression was assessed *via* qPCR. *Gapdh* and *βactin* were used as reference genes. Schematic illustration was created using BioRender.com. Data are means ± S.E. Statistical significance was determined using the one sample t-test. N=3. *p-value<0.0332; **p-value<0.0021; ***p-value<0.0002. **D** RNA of CAFs and CAF*^ΔCD44^* was extracted 48 hours after seeding and subjected to qPCR analysis of expression of *TGFβ1* and *IL10*. *GAPDH* was used as a reference gene. Data are means ± S.E. Statistical significance was determined using the one sample t-test. N=3. *p-value<0.0332; **p-value<0.0021. **E** Immunofluorescence analysis of tumor cell confluency (CellTracker^TM^ Green) in different experimental conditions including or not human T cells (CD45-AF647), CD3xPSMA BiTEs as well as control medium, CAF CM or CAF*^ΔCD44^* CM. Imaging was conducted with the Cell Voyager CQ1 Confocal Quantitative Image Cytometer (Yokogawa) for 24 hours in intervals of 30 minutes. Scale bar, 50 µm. **F** Fitted curves of tumor cell confluency were used to calculate the half-maximal effect (HME) of the two CM conditions. **G** The maximal killing rates at the HME of both conditions were calculated. Statistical significance was determined using a paired, two-tailed t-test. N=4, n=2. ***p-value<0.0002.

### The secretome and immunosuppressive capacity of CAFs is altered after knockout of CD44

By reshaping immune signaling within the TME, CAFs impair T cell activation and contribute to immune evasion. This is mediated by a variety of secreted factors from CAFs that can lead to the recruitment and generation of immunosuppressive cell types [45]. In our study, secretome analyses of CAFs and CAF*^ΔCD44^* revealed significant changes in a variety of secreted factors. The most apparent difference was observed in the secretion of IL-6, interleukin-8 (IL-8), and monocyte chemoattractant protein-1 (MCP-1) (Figure 5A). Correspondingly, the gene expression of both *IL8* and *CCL2* (MCP-1) was markedly reduced (Figure 5B), as observed for *IL6* (Figure 4D).

CAFs were previously shown to corrupt dendritic cells (DC), which are antigen-presenting cells (APC) that induce T cell responses, to become regulatory dendritic cells through the secretion of IL-6 [17]. We therefore investigated how the secreted factors of CAFs and CAF*^ΔCD44^* influence the expression of the immunosuppressive mediators interleukin-10 (IL-10) and indoleamine-2,3-dioxygenase-1 (IDO1) in the LPS-induced dendritic cell line DC2.4. The conditioned medium of CAFs was indeed able to upregulate the expression of *Il10* and *Ido1* in DC2.4 to a higher level compared to the conditioned medium of CAF*^ΔCD44^* (Figure 5C). The same was observed in indirect transwell co-cultures of CAFs or CAF*^ΔCD44^* with DC2.4.

CAF-derived factors have been shown to exert immunosuppressive functions not only on APC but also on T cells directly [45]. Interestingly, two of the most potent immunosuppressive cytokines influencing T cells, TGFβ1 and IL-10, were significantly less expressed after knockout of *CD44* in the CAFs (Figure 5D). To determinate whether CAF*^ΔCD44^* affected CD8^+^ T cells differently compared to CD44-expressing CAFs, we assayed T cell cytotoxicity with the CQ1 confocal live-cell imaging system (Figure 5E [zoomed region from Figure S4A], Figure S4B [Video]). We isolated CD3^+^ T cells from human peripheral blood mononuclear cells (PBMCs) and co-cultured them with endogenously PSMA-expressing LNCAP cancer cells for 24 hours while they were being incubated in the conditioned media of CAFs and CAF*^ΔCD44^* or control medium containing bi-specific T cell engagers (BiTEs) against both CD3 as and PSMA. Staining the tumor cells with CellTracker Green enabled us to assess tumor cell confluency in real time. (Figure S4B). Based on the measured confluency (Figure S4C), the half-maximal effect (HME) was calculated (Figure 5F). T cells that were incubated with the conditioned medium of CD44-expressing CAFs were less efficient in killing the target cells, reflected in the prolonged time to reach the HME (5 hours), contrary to T cells that were incubated with the conditioned medium of CAF*^ΔCD44^*(<4 hours). Additionally, the maximal killing rate at the HME was significantly higher in the CAF*^ΔCD44^* CM-treated T cells compared to the CAF CM-treated T cells (Figure 5G). These results show that CAFs lacking CD44 have a reduced ability to induce/support the immunosuppressive function of immune cells of the tumor microenvironment.

## Discussion

Cancer-associated fibroblasts (CAFs) are a major cellular component of the pancreatic ductal adenocarcinoma (PDAC) microenvironment, yet their functional roles remain incompletely understood. CAFs are known to contribute to key hallmarks of PDAC, including the extensive deposition of desmoplastic extracellular matrix (ECM) and the establishment of a profoundly immunosuppressive tumor microenvironment [46]. However, although therapeutic strategies aimed at targeting CAFs or their interactions with cancer or other stromal cells have shown anti-tumorigenic potential, CAF depletion or broad inhibition can also enhance metastasis and reinforce immunosuppression [22, 47, 48]. The concept of CAF heterogeneity may reconcile these apparently conflicting findings. Besides the two most predominant subtypes, myCAFs and iCAFs, other distinct CAF subtypes have been described, including antigen-presenting CAFs (apCAFs [10]) and interferon-regulated CAFs (ifCAFs [49]), highlighting the functional diversity within this stromal compartment. Nevertheless, myCAFs and iCAFs remain the two most prominent and best-characterized CAF subpopulations in PDAC [50].

In this study we have shown that stromal CD44 is a central regulator of myCAFs and iCAFs in PDAC, thus positioning it as a crucial molecular hub that integrates mechanical and immune signaling within the tumor microenvironment. By combining genetic and functional experiments, we have demonstrated that CD44 expression in fibroblasts is required for their activation into contractile, matrix-producing, and cytokine-secreting CAFs, and that its loss reprograms these cells toward a quiescent and non-activated phenotype. This dual regulation of ECM remodeling and cytokine signaling has revealed a unified mechanism by which CD44 coordinates both myCAF and iCAF properties in PDAC stroma.

Mechanistically, our findings indicate that CD44 acts as a co-receptor in TGFβ1 signaling, a key driver of CAF activation and desmoplasia [51]. In wild-type stellate cells, CD44 facilitates phosphorylation of SMAD2 and associates with TGFβRI upon TGFβ1 induction. Such a physical and functional link has been proposed in metastatic breast cancer cells [52]. There, the binding of HA to CD44 is indirectly involved in the phosphorylation of TGFβRI. However, the TGFβ1-mediated phosphorylation was not influenced, in contrast to our study in imPSCs, where this attenuation of the TGFβ1-mediated Smad signaling prevents the full acquisition of the myCAF phenotype, shown by the reduction of αSMA and collagen I expression. Olsen and colleagues, however, showed that TGFβ1 alone is not sufficient to fully promote myofibroblastic polarization in hepatic stellate cells [53]. Matrix stiffness and its sensing through the YAP/TAZ pathway play a crucial role in fully establishing a myofibroblastic feed forward loop, a process known as mechanosensing. Activated fibroblasts also remodel and stiffen the matrix around them, ensuring their own sustained activation [54].The accompanying changes in CAF morphology, cytoskeletal organization and their capability to contract collagen matrices, as shown in our study suggest that CD44 also contributes to this mechanotransduction, potentially by linking TGFβ1/Smad activity to the YAP/TAZ pathway, which integrates matrix stiffness and actomyosin tension [55–57].

Using a fibroblast-derived 3D matrix system, we evaluated the topography of imPSC-derived matrices and discovered that upon TGFβ1-mediated activation, *Cd44*-depleted imPSCs produce isotropic matrices with fewer organization levels of fibers than *Cd44*-expressing imPSCs. Importantly, we showed fewer parallel fiber organization features that likely promote disease progression through stiffness-regulated mechanotransduction signaling [58, 59]. In our study, we provide evidence that CD44 is important for matrix remodeling that supports pancreatic cancer invasion. In line with our model, Luong & Cukierman demonstrated that pharmacological modulation of TGFβ signaling can normalize pancreatic CAFs by altering ECM alignment and reducing pro-tumorigenic cytokine secretion [60]. In agreement with this model, Alexander and colleagues demonstrated that the f-actin organizing protein palladin is upregulated in response to TGFβ1 and is indispensable for maintaining CAF activation, ECM alignment, and the secretion of immunosuppressive cytokines in PDAC [61]. This suggests that CD44 may act upstream of the TGFβ1-palladin axis, serving as a scaffold that enhances receptor signaling and thereby promotes actin cytoskeletal organization. The loss of CD44 could therefore simultaneously dampen SMAD phosphorylation and palladin-dependent cytoskeletal remodeling, resulting in the isotropic, less fibrotic, and less inflammatory stroma we observed.

The functional consequences of CD44 deletion extend beyond stromal architecture and the myofibroblastic phenotype. The iCAF phenotype is triggered by the exposure to IL-1β resulting in a high expression of cytokines like IL-6 and downregulation of collagen I (Biffi et al., 2019, Figure 4E). In CD44-deficient CAFs, we showed that IL-1β is not able to induce the expression of the inflammatory CAF activation marker *Il6* or the CAF marker *FAP*, suggesting that CD44 is also involved in the IL-1β-mediated induction of the iCAF phenotype. Interestingly, in chondrocytes, knockdown of CD44 attenuated the IL-1β-mediated upregulation of IL-6, supporting the suggested role of CD44 in this pathway [62].

Besides IL-6, other key immunosuppressive cytokines like IL-8, CCL2, IL-10 and TGFβ1 are downregulated in CD44-deficient CAFs. In several cancer types, these factors directly or indirectly affect the T cell-mediated anti-tumor response. IL-6, IL-8 and CCL2 were shown to affect the polarization of macrophages into tumor-promoting M2 TAMs [13], to drive the conversion of dendritic cells into regulatory DCs [17] and to support MDSC differentiation and recruitment [14, 63], all suppressing a functional T cell response against the tumor [18, 45]. Consistent with these findings, our *in vitro* data show that both CD44-deficient pancreatic CAFs and their conditioned medium have a reduced ability to induce a regulatory DC phenotype as shown by diminished expression of *Il10* and *Ido1*. Deletion of CD44 in CAFs also relieved suppression of direct T cell-mediated tumor cell killing by significantly elevating the killing rate of T cells. This might result from the downregulation of CAF-derived TGFβ1 or IL-10 in CAFs that decrease essential cytolytic enzymes in CD8^+^ CTLs [64] or induce the expression of PD-L1 on tumor cells, thus limiting CD8^+^ T cell responses [65].

The dual suppression of TGFβ1- and IL-1β-dependent programs implies that CD44 functions upstream of CAF subtype specification, controlling both myCAF and iCAF trajectories. These results provide direct evidence that fibrosis and immune evasion are mechanistically coupled through CD44.

Therapeutically, depletion of CD44 in CAFs offers a strategy to remodel the desmoplastic and immune-excluded landscape of PDAC into a more responsive and treatable state. In contrast to CAFs-depletion studies, our data present a more nuanced strategy for functional reprogramming of CAFs *via* CD44 inhibition to normalize rather than eliminate the stroma.

## Methods

### Mice

B6.Cg-Tg(Pdgfrb-cre/ERT2)6096Rha/J mice [31] were crossed with *C57BL/6-Tg(Cd44-[exon3]^flox/flox^* mice to generate the *Cd44^flox/flox^Pdgfrb-CreER^T2^* (*Cd44^fl/fl^;PdgfrβCreER^T2^*) mouse line. The resulting line was used for CAF-specific *Cd44* knockout experiments. For *in vivo* purposes, only male mice were subjected to experiments. Animals were assigned to experimental groups regarding their genotype. B6.Cg-Tg(Gfap-cre)73.12Mvs/J [66] were crossed with *C57BL/6-Tg(Cd44-[exon3]^flox/flox^* mice to generate the *Cd44^flox/flox^Gfap-Cre* (*Cd44^fl/fl^;GfapCre*) line. For pancreatic stellate cell isolations, only male mice were used, and animals were selected based on their genotype. All animals were housed and maintained in facilities approved by the Regierungspräsidium Karlsruhe (Germany) under specific pathogen-free conditions and were handled according to EU directives for animal experimentation. The experiments were authorized by the Regierungspräsidium (35-9185.81/G-10/19) and termination criteria were approved. They were defined to preclude suffering, unnecessary pain and harm to the animals while enabling accurate and timely data generation.

### Cell lines

FC1245 cells were kindly gifted by Dr. Dave Tuveson (Cold Spring Harbor Laboratory, Cold Spring Harbor, NY). HEK293T cells were obtained from ATCC (Wesel, Germany) and imPSCs were obtained from Dr. Angela Mathison and Dr. Raul Urrutia (Medical College of Wisconsin, Milwaukee, USA) [32]. These cell lines were cultured in DMEM containing GlutaMAX supplement (Gibco) and 10% fetal bovine serum (FBS, Gibco) and 1% Pen/Strep (P/S, Gibco). Human pancreatic CAFs were obtained from Vitro Biopharma (Denver, CO, USA) and were cultured in MSC-GRO™ Pancreatic CAF Maintenance Medium (Vitro Biopharma, Inc.) with 1% P/S (Gibco). DC2.4 cells were obtained from Merck and were cultured in RPMI (10% FBS, 1% GlutaMAX supplement, 10% 1 M HEPES, 0.1% β-mercaptoethanol, 1% non-essential amino acids, 1% P/S). Cell lines were tested regularly for mycoplasm contamination.

### Orthotopic PDAC cell implantation

Six- to ten-week-old *Cd44^fl/fl^;PdgfrβCreER^T2^*or control mice were injected intraperitoneally with 20 mg/ml of Tamoxifen dissolved in peanut oil, once per day for 5 days. At day seven, 4.5 x 10^4^/ 30 µl FC1245 cells in sterile PBS (Gibco) were injected. For this, isoflurane was used as an inhalation anesthetic and was vaporized with compressed air at 1 liter per minute. The abdominal skin of the animals was cleaned with 70 % ethanol. A 5 mm incision was made at the left abdominal flank with sterile micro-scissors. The spleen and part of the pancreas were gently pulled through the incision with blunt forceps. 30 µl of cell suspension were injected into the pancreas through a 1-ml syringe with a 30G of ½ -inch needle, 3 mm medial to the spleen hilium. The pancreas was inspected for 1 min for possible hemorrhage and leakage, before it was, together with the spleen, replaced into the peritoneal cavity with blunt forceps. Subsequently, muscle and skin layers were closed with interrupted 4-0 non-reabsorbable sutures. The volume was calculated using the following formula: volume = length x width x height x 0.5 (mm^3^).

### Isolation of fibroblasts from the pancreas of Cd44^fl/fl^;PdgfrβCreER^T2^ mice

Fibroblasts from the pancreas of *Cd44^fl/fl^;PdgfrβCreER^T2^*mice were isolated using a protocol modified from Waise and colleagues [67]. Mice were sacrificed and the pancreas was excised. Single-cell suspensions were established using the Miltenyi MACS® mouse tumor dissociation kit (Miltenyi Biotec) according to the manufacturer’s protocol. After collection of the single cells by centrifugation for 7 minutes at 300g, they were washed with ammonium-chloride-potassium (ACK) to lyse red blood cells. Cells were then collected *via* centrifugation at 300g for 7 minutes and were resuspended in DMEM containing GlutaMAX supplement (Gibco) and 10% fetal bovine serum (FBS, Gibco) and 1% Pen/Strep (P/S, Gibco) and seeded on culture plates. After attachment, the plates were washed three times with PBS before adding DMEM (GlutaMAX, 10% FBS, 1% P/S) to allow cells to grow.

### 3D culture of CAFs

5x10^3^ CAFs or CAF*^ΔCD44^* were seeded in 50 µl Matrigel (Corning) domes into 24-well plates and cultured in 500 µl of MSC-GRO™ pancreatic CAF maintenance medium (Vitro Biopharma) with 1% of P/S. Retrieval of cells from the Matrigel was achieved by incubation with 250 µl of pre-warmed dispase (Stemcell Technologies,1 U/ml). Collected cell pellets were prepared for further analysis.

### Cell culture

For immunofluorescence analyses, 1x10^4^ imPSCs/imPSC*^ΔCD44^*or CAFs/CAF*^ΔCD44^* per well were seeded on glass slides in 12-well plates. The cells were induced with 50 ng/ml recombinant TGFβ1 (PeproTech) in DMEM containing 1% FBS and 1% P/S for 24 hours.

For induction experiments, cells were either seeded in Matrigel (see 3D culture of CAFs) or in flasks or plates. In 2D culture, CAFs/CAF*^ΔCD44^*(1x10^5^) or imPSCs/imPSC*^ΔCD44^* (9x10^4^) were seeded into 6-well plates overnight. Cells were starved in DMEM containing 1% FBS and 1% Pen/Strep for 24 hours, before cells were induced with 50 ng/ml of recombinant TGFβ1 (either 24 or 72 hours) or 20 ng/ml of IL-1β or IL-6 (PeproTech). Cells were either prepared for protein or RNA analyses or removed from the Matrigel for RNA isolation.

### Cd44 knockdown using siRNA

9x10^4^ imPSCs were seeded into 6-well plates. 100 pmol of *Cd44*-specific siRNA or non-silencing control siRNA were diluted in 100 µl of DMEM (without FBS) per well and incubated with 12 µl of HiPerFect (Qiagen) for 7.5 min at RT. The complexes were added dropwise onto the cells and incubated. After 24 hours, a second transfection was performed, and cells were seeded for the respective experiments after 48 hours of total transfection. The sequences will be provided upon request.

### Lentivirus production and cell transduction

The CRISPR/Cas9-mediated knockout of *Cd44/CD44* in imPSCs and CAFs was performed using a third-generation lentiviral system. HEK293T packaging cells were transfected with pMDL (*gag* and *pol* genes), REV (*rev* gene) and VSV-G (*vsv-g* gene). Depending on the approach the fourth plasmid either encoded a Cas9 endonuclease and a small guide RNA (sgRNA) to target the *Cd44* gene sequence (mouse: VB180221-1067cyf human:VB180221-1056peh) or a scramble sequence tagged with mCherry (VB240604-1056nya).

The plasmids were transfected into HEK293T cells using PromoFectin (PromoCell) according to the manufactureŕs protocol (5 µg of PMDL, 2.5 µg of REV, 2.8 µg of VSV-G and 10 µg of the expression vector). After six hours of transfection, the medium was removed, and 5.5 ml of growth medium were added into the culture plate. After 24 hours of particle production, the medium was removed and filtered through a 0.45 µm sterile filter on top of the target cells. This was repeated once. For selection, the cells were treated with the nucleoside antibiotic puromycin (InvivoGen, 2 µg/ml) and analyzed and sorted for CD44-negative or mCherry-positive cells (scramble) via flow cytometry.

### Flow cytometry

CRISPR/Cas9-transduced CAFs or imPSCs were collected, incubated with human (Clone FC1, BD Pharmingen^TM^ RRID: AB_2728082) or mouse Fc Block^TM^ (Purified rat anti-mouse CD16/CD32; BD Pharmingen^TM^; RRID: AB_394656) for 30 minutes and stained with a PE anti-mouse/human CD44 antibody (1:100, clone: IM7, BioLegend, RRID: AB_312959) or isotype control (1:100, BioLegend, RRID: AB_326552) for 15 minutes. Cells were washed with FACS buffer (PBS + 2 % FBS + 2 mM EDTA) and analyzed by flow cytometry using a FACSAria Fusion cytometer (BD Biosciences).

### Indirect immunofluorescence

Cells on coverslips were fixed with 2% paraformaldehyde in PBS for 15 minutes at RT, washed with PBS and permeabilized with 0.1% Triton-X for 15 minutes at RT. After incubation in blocking buffer (5% FBS in PBS) for 45 minutes at RT, samples were incubated with primary antibodies against CD44 (IM7, 2.5 µg/ml, BD Biosciences, RRID: AB_393732), COL1A1 (1:200, Cell Signaling, RRID: AB_2800169), Phalloidin (1:1000, Alexa Fluor 488, Thermo-Fisher Scientific), GFAP (1:200, Cell Signaling, RRID: AB_2631098) or ɑSMA (1:200, Cell Signaling, RRID: AB_2857972) diluted in 5% FBS in PBS overnight at 4°C. Control samples were incubated with blocking buffer only. After washing with PBS-T, cells were incubated with the secondary antibodies goat anti-rat IgG secondary antibody, Alexa Fluor® 488 conjugate (1:1000; Thermo Fisher, RRID: AB_2534074, goat anti-rabbit IgG secondary antibody, Alexa Fluor® 546 conjugate (1:1000; Thermo Fisher, RRID: AB_2534115) and DAPI (1:1000, Sigma-Aldrich) diluted in 5% FBS in PBS at RT for 30 minutes. Coverslips were mounted onto microscope slides using Mowiol 4-88 (Roth) mounting medium and dried overnight. The slides were stored at 4°C until analysis with a Zeiss LSM 800 confocal microscope. Second harmonic generation signal was acquired at 430 nm by using linear unmixing mode after excitation at 860 nm.

### RNA extraction, reverse transcription and qPCR

Total RNA was extracted using the RNeasy Mini Kit (Qiagen). For the generation of cDNA from freshly isolated mRNA the qScriber^TM^ cDNA Synthesis (highQu) was used according to the manufacturer’s protocol. The cDNA was then diluted in a 1:4 ratio in RNase free water and the qPCR reactions (20 µl) consisted of 10 µl 2x GoTaq® qPCR Master Mix (Promega), 0.5 mM reverse and forward primers, and 2 µl cDNA. The qPCR was performed according on the CFX96 Touch Real-Time PCR detection system (Biorad). *Gapdh* and *βactin* were used as reference genes and the relative quantification was calculated based on the Litvak method (ΔΔCt). Primers were designed using the Primer3 open software (http://primer3.ut.ee) using transcript sequences from the NCBI-database.

### Co-cultures

#### Incubation of DC2.4 cells with CAF/CAF^ΔCD44^-conditioned medium (CM)

2 x 10^5^ DC2.4 cells were seeded in a 6-well plate and incubated with their full RPMI-1640 medium or with RPMI-1640 medium containing 50% of either CAF or CAF^ΔCD44^ CM. At 24 hours after seeding, 1 µg/ml LPS was added and incubated for 24 hours before RNA was extracted for downstream analysis.

#### Indirect Co-culture of CAFs/CAF^ΔCD44^ with DC2.4 Cells

DC2.4 cells were seeded in a 12-well plate with 1 x 10^5^ cells per well. 5 x 10^4^ CAFs/CAF^ΔCD44^ were seeded in ThinCert inserts (0.4 μm pore, Greiner Bio-One) on top of the DC2.4. After 24 hours of co-culture, 1 µg/ml of LPS were added to the well and incubated for 24 hours, before RNA was extracted from the DC2.4 for downstream analysis.

### Protein detection

#### Human cytokine array

Secretome analysis of CAFs/CAF*^ΔCD44^* were conducted using the Human Cytokine Antibody Array (Abcam) according to the manufacturer’s instructions. 2x10^5^ CAFs/CAF*^ΔCD44^*were seeded and incubated for 48 hours. Media was collected, centrifuged (450 rcf, 6 min) and filtered through sterile 0.2 µm syringe filters. The array membranes were incubated with conditioned media and through a biotin-streptavidin-HRP antibody system, the factors present were visualized with the ChemiDoc™ Touch Imaging system (BioRad) upon administration of the HRP substrate. Positive and negative control spots were used to quantify the relative signal intensity.

### Cell-derived matrices production

The procedure was adapted from Kaukonen et al., *Nat Protoc*, 2017, with minor modifications [38]. Plates with sterile coverslips were coated with 1% gelatin and crosslinked with 1% glutaraldehyde (v/v) for 20 min at RT. Crosslinking was quenched with 1M glycine for 20 min at RT, followed by PBS washes. Cells were seeded at 0.4x10⁶ per well in DMEM supplemented with 10% FBS and 1% antibiotics. At 48h hours post-seeding, extracellular matrix deposition was stimulated by daily treatment with 1% ascorbic acid (50μg/mL) with or without TGFβ1 (5ng/ml) for 7 days. Fibroblasts were then removed using extraction buffer (100mM NH4OH in PBS containing 0.5% (v/v) Triton X-100) for 3 min, followed by two PBS washes. Remaining DNA was digested with DNase I (10 µg/mL) for 2x30 min at 37°C. Cell-derived matrices were stored at 4°C in PBS or fixed with 4% paraformaldehyde for 10 min prior to fluorescent staining and confocal microscopy.

### Contraction assay

The contraction assay was adapted from Chitty et al., *Cancer Reports*, 2020, with minor modifications [68]. 96-well plates were coated with 2% BSA and 2.5x10⁴ cells were embedded in 100 µL of rat tail collagen I hydrogel (1.5 mg/mL, Corning). Hydrogels were polymerized for 1h at 37°C, after which 100 µL of complete growth medium was added to each well. Gels were allowed to contract for 48h at 37°C. Images of the collagen lattices were acquired using a MICA microscope (Leica), and the diameter of the well and gel was measured using ImageJ.

### T cell killing assay

CD3^+^ T cells were isolated from human PBMCs using the Pan T cell isolation kit (Miltenyi Biotec, 130-096-535) according to the manufacturer’s protocol. The T cells were activated for three days using CD3/CD28 activation beads (Dynabeads^TM^ Human T-activator CD3/CD28, Gibco^TM^) at a 1:1 bead-to-cell ratio and 30 U/mL recombinant IL-2 (R & D Systems, BT-002-050) in serum-free X-VIVO 15 medium (Lonza). Target tumor cells were seeded 20 hours before the start of the experiment on fibronectin-coated 96-well imaging plates (Screenstar Microplate, Greiner Bio-One; fibronectin: R&D Systems 1030-FN-05M). On the day of co-culture tumor cells were stained with CellTracker^TM^ Green CMFDA (Invitrogen, Thermo Fisher). T cells and target cells were suspended in the experimental (60 % CAF conditioned media and 40 % RPMI-1640 (HiGlutaXL RPMI-1640, HiMedia, AL028G) + 10 % FBS (Bio & Sell, FBS.S0615)) and control media containing Anti-CD45-AF647 [1:500, Invitrogen^TM^, Thermo Fisher, MA5-38730] for labelling T cells and NucSpot 568/580 [1:2000, biotium, 41036] for identifying dead cells. The CD3xPSMA BiTEs (MedChemExpress, HY-P99802) were also prepared in the respective experimental and control medium and added to the co-culture at 10 µg/mL. Imaging was conducted with the Cell Voyager CQ1 Confocal Quantitative Image Cytometer (Yokogawa) for 24 hours at intervals of 30 minutes. CellTracker Green was recorded with excitation at 488 nm with 20 % laser power and 500 ms exposure time using a BP525/50 emission filter. NucSpot 568/580 was recorded with excitation at 561 nm at 20 % laser power and 500 ms exposure time using a BP617/73 emission filter. CD45-AF647 was recorded with excitation at 640 nm at 20 % laser power and 500 ms exposure time using a BP685/40 emission filter. Maximum intensity projection images were recorded from three Z planes covering a 3 µm using a 20x/0.8 NA objective (Olympus, UPLXAPO20X). Imaging was performed with four biological replicates of each experimental media in technical duplicates per media condition.

The tumor cell confluency was measured based on the green CellTracker fluorescence using CellPathfinder (version 3.06.01.08, Yokogawa). The confluency over time was baseline-corrected and depicted as percent difference [100*(Value-Baseline)/Baseline]. The technical replicates were treated as repeated measures.

The resulting curves were fitted using the [Inhibitor] vs. response -- Variable slope four parameter logistic models [Y=Bottom + (Top-Bottom)/(1+(IC50/X)^HillSlope)]. To calculate the maximal killing rate, the first derivative at t=half-maximal effect (HME) was calculated using:

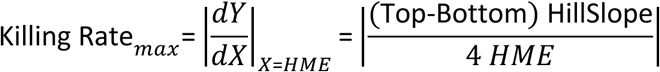

The resulting maximal killing rates were compared between the two conditioned media conditions. Significance was analyzed using a paired, two-tailed t-test. All analyses were performed using GraphPad Prism (version 10.3.1, GraphPad Software, Boston, Massachusetts USA) and Microsoft Excel (Microsoft 365 MSO, Version 2508 Build 16.0.19127.20314, Microsoft, USA)

### Single cell data analysis

Raw read counts were obtained from the Genome Sequence Archive (GSA; CRA001160). Data processing and quality control were performed using Scanpy. Low-quality cells were filtered out according to the following criteria: only cells expressing between 100 and 6,000 genes and with fewer than 100,000 total counts were retained. Genes detected in fewer than three cells were excluded. Doublet detection was performed using the scanpy.pp.scrublet function, identifying 0.26% of cells as potential doublets, which were excluded from downstream analyses. After filtering, 45,776 cells remained for further analysis.

Read counts were normalized per cell to a total count of 10,000. Principal component analysis (PCA) was performed on the 5,000 most variable genes, followed by nearest-neighbor graph construction using the top 30 principal components. UMAP was applied for dimensionality reduction and visualization. Cell type annotations from the original publication were used as the reference labels. Marker gene analysis was conducted using the Wilcoxon rank-sum test, comparing each cell type against all others. Comparisons between pancreatic cancer and normal conditions were performed using the same approach. Genes with an adjusted p-value below 0.05 were considered as significantly regulated. Fibroblastic cells were defined by the expression of ɑSMA, PDGFRβ, lumican (LUM), decorin (DCN), adipognesis regulatory factor (ADIRF), COL1A1.

### Statistical analysis

Tests for Gaussian (normal) distribution were conducted for all data by Shapiro-Wilk normality test or Kolmogorov-Smirnov normality test. Upon passing the normality test, both samples were assumed to derive from populations with the same variances. Using a non-paired parametric tests like the one-sided unpaired Student’s t-test, mean values of quantitative variables between two independent groups were compared. In case of comparison between the mean of a single sample against a hypothetical mean a one-sample t-test was used. If means of more than two samples were compared, a one-way analysis of variances (ANOVA) was performed and Holm-Šídák’s multiple comparison was used as a post-hoc test. Data are shown as the average with ± standard error of mean (SEM). We accepted a significance level α <0.05. The p-value is indicated by asterisks and defined as * = 0.032, ** = 0.0021, *** = 0.00021, **** < 0.0001. Statistical analysis was performed using GraphPad Prism 9.3.1 software (GraphPad, RRID:SCR_002798).

## Acknowledgements

SMT, YMH, JM, SJS, LMM, LMS, MC and VO-R were supported by the Helmholtz program “Materials Systems Engineering (MSE)”. We thank the Deutsche Forschungsgemeinschaft (DFG, German Research Foundation) for the support: OR 124/24-1. We acknowledge funding from the German Federal Ministry of Education and Research (BMBF) within the Medical Informatics Funding Scheme EkoEstMed–FKZ 01ZZ2015 (G.A.). We thank the animal facility at the IBCS-FMS. We thank Prof. Dave Tuveson (Cold Spring Harbor Laboratory, Cold Spring Harbor, NY) for providing the FC1245 cell line. We thank Associate Professor Angela Mathison and Prof. Raul Urrutia (Medical College of Wisconsin, Milwaukee, USA) for providing the imPSC cell line [32]. We thank Dr. Elvire Guiot (Imaging Center (ICI), IGBMC, Strasbourg) for the SHG analysis. We thank Prof. Ulrich Rothbauer (Eberhard Karls University, Tübingen) for providing the Vimentin chromobodies.

## Funding Declaration

Helmholtz program “Materials Systems Engineering (MSE)”: SMT, YMH, JM, SJS, LMM, LMS, MC and VOR.

Funded by the Deutsche Forschungsgemeinschaft (DFG, German Research Foundation)-OR124/24-1.

German Federal Ministry of Education and Research (BMBF) within the Medical Informatics Funding Scheme EkoEstMed–FKZ 01ZZ2015: GA.

TRON: LL, ES.

## Author contribution

SMT, YMH, JM: Performance of experiments, analysis, preparation of figures. SMT, YMH, JM and VOR: Preparation of the manuscript. VOR: Conceptualizing the project, acquisition of funding, providing resources and supervised the project. LL, ES, LMM, SJS and MC: Performance of experiments. GA: Analysis of publicly available RNA Sequencing data. LMS: Design of lentiviral vectors. All authors read and approved the manuscript. SMT and YMH are shared first authors.

## Data availability

RNA sequencing data from Peng et al. (2019) [29]: The accession number for the sequencing data reported in this paper is GSA: CRA001160. These data have been deposited in the Genome Sequence Archive under project PRJCA001063.

