## Supplementary data for "Loss of CD44 re-educates pancreatic cancer-associated fibroblasts modulating their fibrotic and immunosuppressive functions"

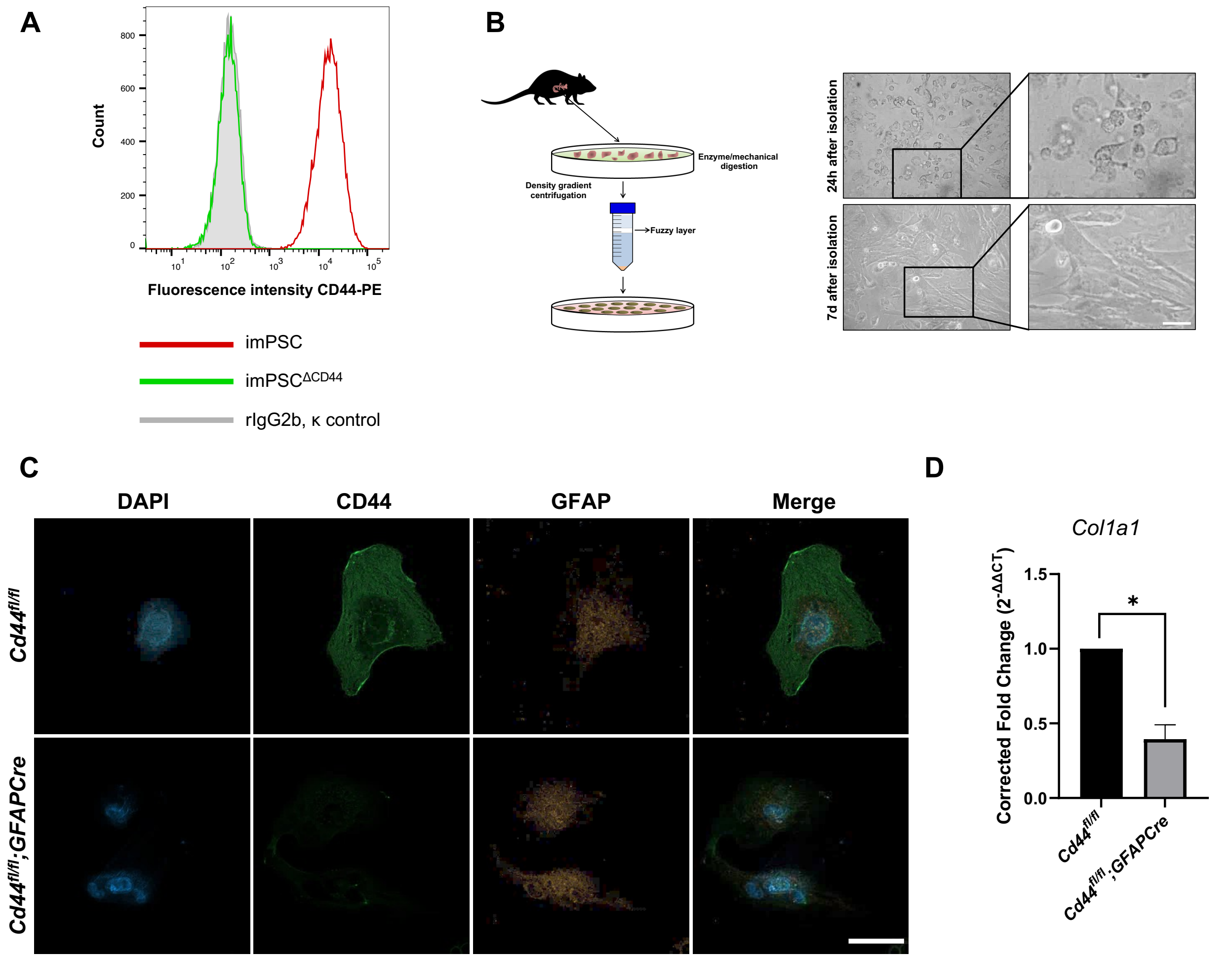


**Supplementary Figure 1 | *Cd44* knockout in CAFs and PSCs in *Cd44*-floxed mouse models.** **A** Flow cytometry analysis of the CD44 expression in imPSCs after CRISPR/Cas9-mediated knockout. imPSCs and imPSC$^{\Delta CD44}$ cells were stained with panCD44 antibodies (IM7) coupled to phycoerythrin (PE) and the corresponding rat IgG2b, κ isotype control, also labelled with PE. **B** Isolation of PSCs from healthy pancreatic tissue using Nycodenz gradient centrifugation. Brightfield images show PSCs 24 hours and 7 days after isolation. **C** Immunofluorescence analysis of freshly isolated PSCs from *Cd44*$^{fl/fl}$*;GfapCre* and corresponding control mice stained for CD44 (AlexaFluor488) and glial fibrillary acidic protein (GFAP, AlexaFluor546). **D** Freshly isolated PSCs from *Cd44*$^{fl/fl}$*;GfapCre* and control mice were treated for 24 hours with recombinant TGFβ1. Expression of *Col1a1* was analyzed by qPCR. *Gapdh* and *βactin* were used as reference genes. Data are means ± S.E. Statistical significance was determined using the one sample t-test. N=3. *p-value<0.0332

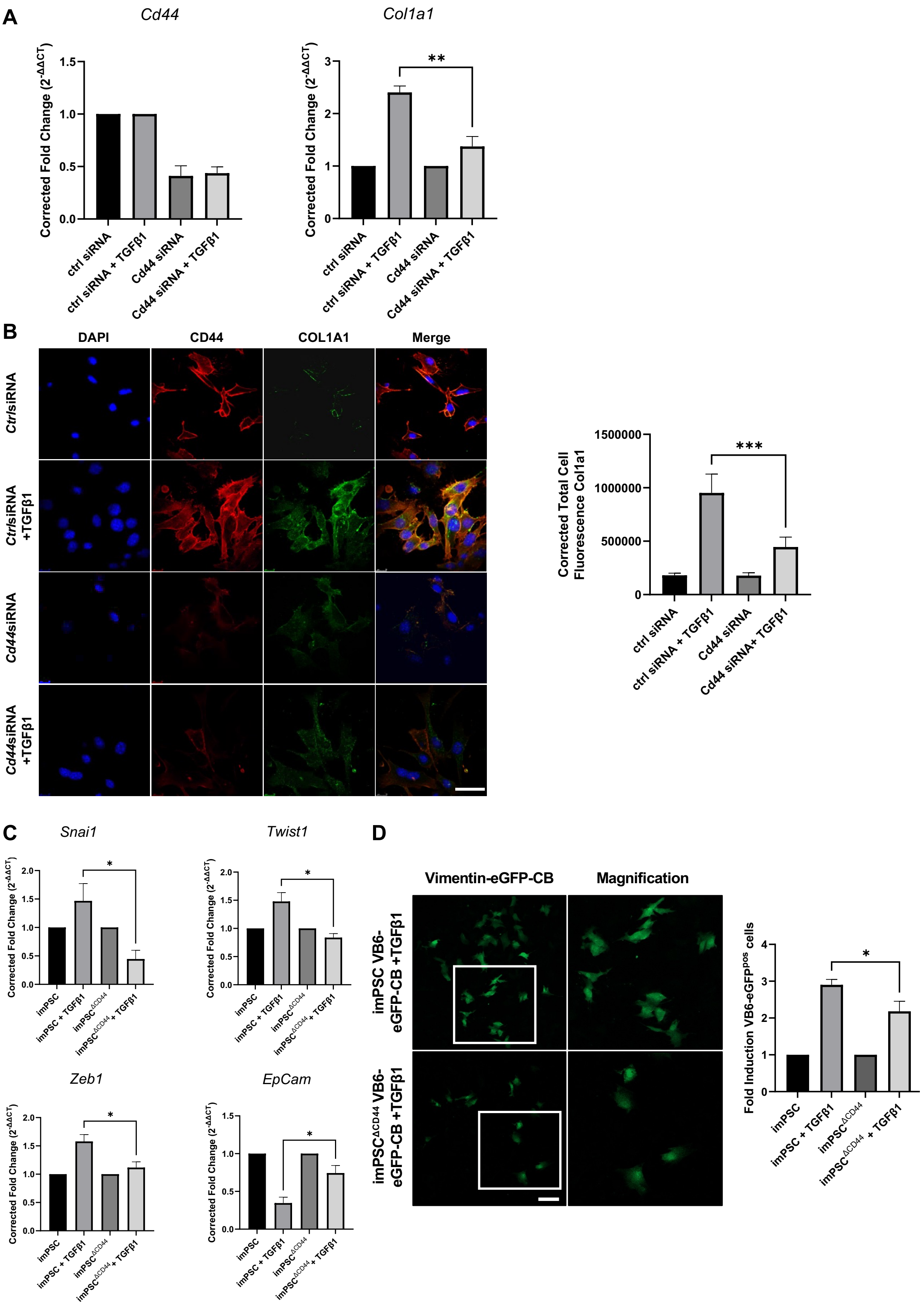
A
Cd44
Col1a1
Corrected Fold Change (2^-ΔΔCT)
1.5
1.0
0.5
0.0
3
2
1
0
**
ctrl siRNA
ctrl siRNA + TGFβ1
Cd44 siRNA
Cd44 siRNA + TGFβ1
B
DAPI
CD44
COL1A1
Merge
Ctrl siRNA
Ctrl siRNA +TGFβ1
Cd44siRNA
Cd44siRNA +TGFβ1
Corrected Total Cell Fluorescence Col1a1
1500000
1000000
500000
0
***
ctrl siRNA
ctrl siRNA + TGFβ1
Cd44 siRNA
Cd44 siRNA+ TGFβ1
C
Snai1
Twist1
Zeb1
EpCam
Corrected Fold Change (2^-ΔΔCT)
2.0
1.5
1.0
0.5
0.0
*
imPSC
imPSC + TGFβ1
imPSC^ΔCD44
imPSC^ΔCD44 + TGFβ1
D
Vimentin-eGFP-CB
Magnification
imPSC VB6-eGFP-CB +TGFβ1
imPSC^ΔCD44 VB6-eGFP-CB +TGFβ1
Fold Induction VB6-eGFP^pos cells
4
3
2
1
0
*
imPSC
imPSC + TGFβ1
imPSC^ΔCD44
imPSC^ΔCD44 + TGFβ1

16

**Supplementary Figure 2 | *Cd44* knockout and siRNA-mediated reduction in imPSCs influence activation and EMT. A** ImPSCs were transfected with control siRNA (*Ctrl*siRNA) or siRNA targeting all CD44 isoforms (*Cd44*siRNA) and induced with recombinant TGFβ1 for 24 hours. qPCR analysis was used to determine the expression of *Cd44* and *Col1a1*. *Gapdh* and β*actin* were used as reference genes. **B** Transfected imPSCs were analyzed *via* IF using specific antibodies against CD44 (IM7, AlexaFluor546) and COL1A1 (AlexaFluor488). Nuclei were counterstained with DAPI. Scale bar, 50 µm. The CTCF of all conditions was determined by quantifying the fluorescence *via* ImageJ with deduction of background signals. **C** Expression analysis of *Snai1*, *Twist1*, *Zeb1* and *EpCam* via qPCR after TGFβ1 treatment. *Gapdh* and *βactin* served as reference genes. **D** Chromobody (CB) plasmids encoding the Alpaca-derived vimentin-specific $V_H$H-eGFP fusion protein (VB6-eGFP) were transfected into imPSCs and imPSC$^{\Delta CD44}$ target cells. Representative confocal images show chromobodies binding to endogenous vimentin (green). Scale bar, 50 µm. eGFP intensity upon TGFβ1 stimulation for 72 hours was quantified *via* flow cytometry in imPSCs and imPSC$^{\Delta CD44}$. Data are means ± S.E. Statistical significance of all experiments depicted here was determined using the one-sided unpaired Student's t-test. N=3. *p-value<0.0332, **p-value<0.0021, ***p-value<0.0002.

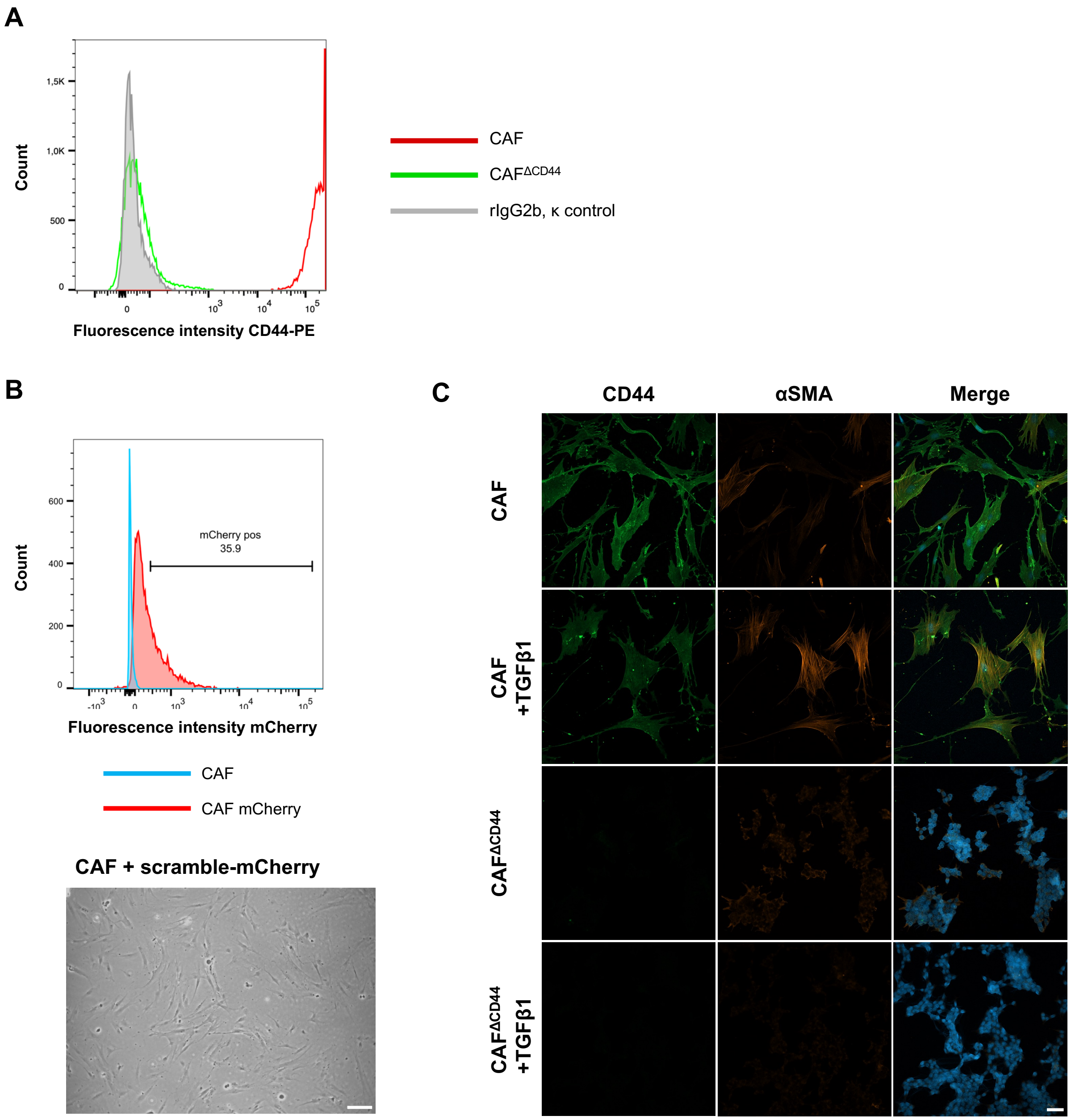


**Supplementary Figure 3 | Generation of CRISPR/Cas9-edited human CAF cell line and corresponding scramble cell line. A** Flow cytometry analysis of the CD44 expression in CAFs after CRISPR/Cas9-mediated knockout after sorting. CAFs and CAF$^{\Delta CD44}$ cells were stained with CD44 antibodies (IM7) coupled to phycoerythrin (PE) and the corresponding rat IgG2b, κ isotype control, also labelled with PE. **B** Flow cytometry analysis of mCherry signal in CAFs and CAFs transduced with a CRISPR scramble construct marked with mCherry. Brightfield picture of CAF transduced with the sgRNA-mCherry. Scale bar, 100 µm. **C** IF analysis of CAFs and CAF$^{\Delta CD44}$

using specific antibodies against CD44 (IM7, AlexaFluor488) and αSMA (AlexaFluor546). Nuclei were counterstained with DAPI. Scale bar, 50 μm. N=1.

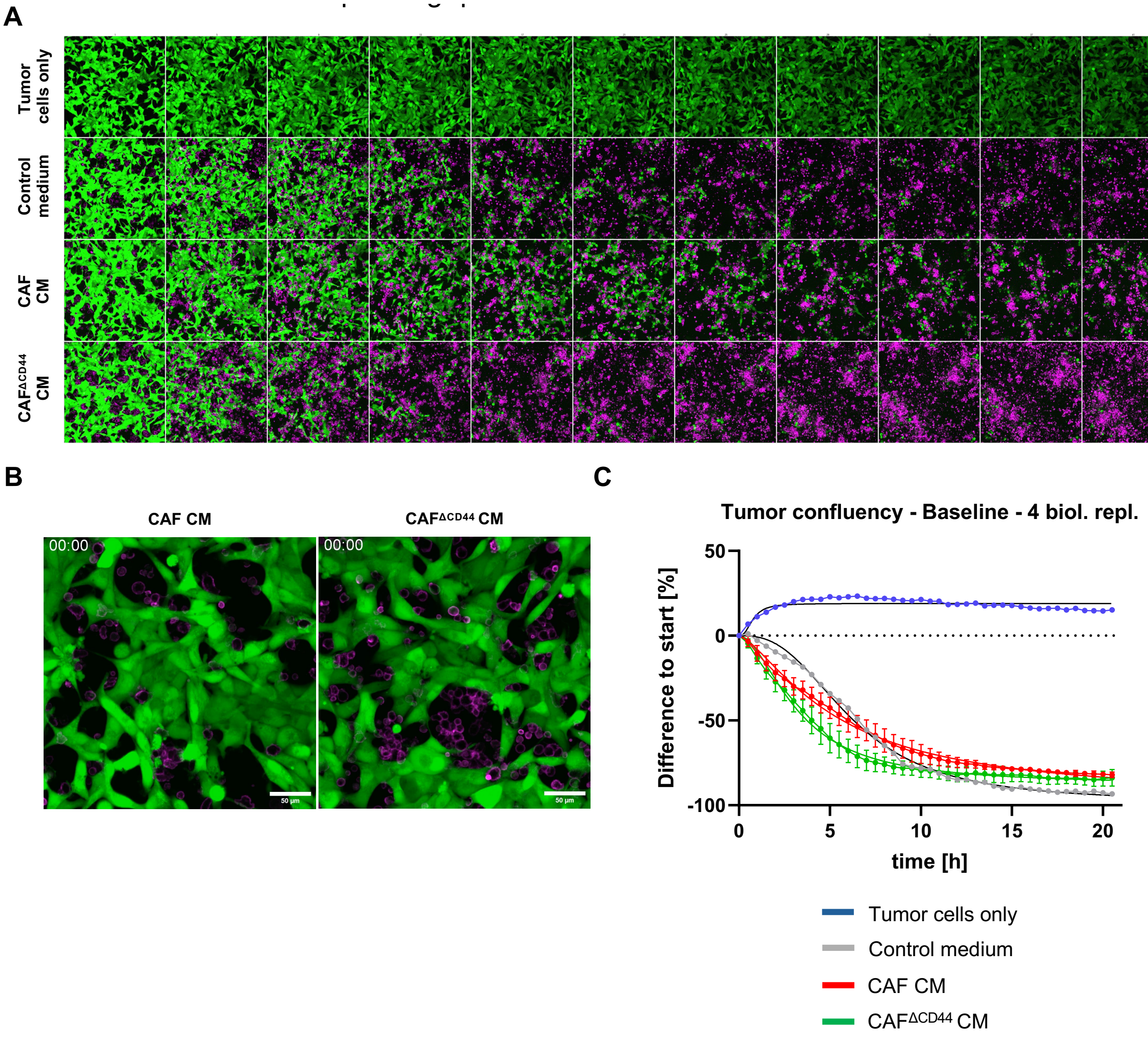


**Supplementary Figure 4 | T cell-mediated tumor cell killing assay in various experimental conditions and corresponding quantifications. A** Overview IF pictures of tumor cell confluency (CellTracker™ Green) and human T cells (CD45-AF647) in all experimental conditions (tumor cells only, control medium, CAF CM or CAF$^{\Delta CD44}$ CM). Images were created in a time span of 24 hours with the Cell Voyager CQ1 Confocal Quantitative Image Cytometer (Yokogawa) at intervals of 30 minutes. **B** Corresponding videos of the CAF CM and CAF$^{\Delta CD44}$ CM-incubated conditions showing T-cell mediated tumor cell killing. **C** The tumor cell confluency was measured based on the green CellTracker fluorescence using CellPathfinder. The confluency over time was baseline-corrected and depicted as percent difference [100*(Value-Baseline)/Baseline] and the curves were fitted. N=4, n=2.

**Supplementary methods:**

**Isolation of quiescent PSCs**

Quiescent PSCs were isolated from the pancreas of *Cd44*$^{fl/fl}$*;GfapCre* following the protocol of Apte *et al.* (1998) with minor modifications. After cervical dislocation, the pancreas was excised, cleared of connective and fatty tissue, and washed in PBS+ 1% penicillin/streptomycin. The tissue was minced with surgical scissors and digested with an enzyme mix (GBSS [Gey's balanced salt solution] with collagenase [1.3 mg/ml], protease [1 mg/ml], DNase [0.01 mg/ml]) at 37 °C for 30 min in a shaking water bath. The suspension was filtered through a 100 µm strainer, washed with cold GBSS containing BSA, and centrifuged (500 g for 10 min at 4 °C). After a second wash, the pellet was resuspended in 9.5 ml cold GBSS + BSA and 8 ml GBSS + Nycodenz (28.7%), divided into two 15 ml tubes, and overlaid with 3 ml GBSS + BSA to form a gradient. Following centrifugation (1400 g for 20 min at 4 °C; lowest acceleration, no brake), the PSC-enriched band was collected, pooled, washed, and centrifuged (500 g, 10 min, 4 °C). Cells were resuspended in complete culture medium and seeded in appropriate vessels for downstream experiments.

**Vimentin-Chromobodies**

VB6-eGFP-Chromobody expression vectors were kindly provided by Dr. Ulrich Rothbauer (NMI, Reutlingen, Germany). imPSC and imPSC$^{\Delta CD44}$ were transfected with the expression vector encoding the VB6-eGFP-Chromobody (CB) sequence [1] using Lipofectamine 3000 (Thermo-Fisher Scientific) according to the manufacturer's protocol. 24 hours post transfection, cells were selected using 0.5 mg/ml G418 for 72 hours. Vimentin expression was induced in selected cells using 50 ng/ml TGFβ1 for 72 hours and eGFP-positive cells were sorted using a FACSAria Fusion (BD Biosciences). For EMT induction experiments, 9x10$^4$ cells were seeded into 6-well plates and treated with 50 ng/ml TGFβ1 in DMEM with 1% FBS and 1% Pen/Strep for 72 hours. Single cell suspensions were analyzed for GFP-positive cells using a FACSAria Fusion (BD Biosciences).
ng/ml TGFβ1 for 72 h, and GFP$^+$ cells were subsequently enriched by flow cytometry.

**Protein isolation and detection**

Cells were lysed with 1% Triton lysis buffer (1 % Triton-X-100; 150 mM NaCl; 50 mM Tris-HCl (pH 7.4); 5 mM $Na_3VO_4$; 25 mM NaF; 0.1 % NP-40; 1 mM EDTA; 1 mM EGTA; 50X protein inhibitor cocktail (Roche); adjust pH to 7.0). Lysates were collected using a rubber scraper and were centrifuged (13 200 rpm, 20 min, 4 °C), followed by the transfer of the supernatant to fresh tubes. 4x Laemmli sample buffer (BioRad) with 10% β-mercaptoethanol was added, and samples were boiled at 95 °C for 5 min.

Co-Immunoprecipitation (Co-IP)

imPSCs were lysed using 1% Triton X-100 buffer. 50 µl of the supernatant was taken as input control and mixed with 4x Laemmli sample buffer (BioRad) with 10% β-mercaptoethanol, heated at 99°C for 5 minutes and stored at -20°C. The remaining supernatant was used for immunoprecipitation. To target the protein of interest and its interacting partners, samples were incubated overnight with the panCD44 (clone: KM201) antibody on an orbital shaker. On the following day, Agarose G beads were washed three times with 1% Triton-X lysis buffer followed by a centrifugation at 1200 rpm for 3 minutes (pre-clearing). Subsequently 30 µl of the beads were added to the samples. For the IgG control, directly coupled beads were used. The samples were incubated for 3h at 4°C on an orbital shaker. After the incubation period, samples were washed three time and were centrifuged (1200 rpm, 3 min, 4°C) to spin down the beads coupled to the antibodies and the proteins. The supernatant was removed, and the beads were resuspended in 1% Triton-X lysis buffer and washed three times for 5 minutes at 4°C on an orbital shaker. After another centrifugation (1200 rpm, 3 min, 4°C), the supernatant was completely aspirated and the proteins were denatured in 2x Laemmli sample buffer (BioRad) with 10% β-mercaptoethanol by boiling at 99°C for 5 minutes.

Western Blot

SDS–PAGE was conducted using 5% stacking and 8% separating gels to separate proteins. Electrophoresis ran for 60 min at 100 V and for 45 min at 120 V. Transfer of proteins onto PVDF membranes was executed with the Trans-Blot Turbo system (Bio-Rad). After blocking of the membranes with 5% BSA in TBS-T for 1 h, they were incubated overnight at 4 °C with primary antibodies (1:200 - 1:1000 in 5% BSA/TBS-T). Membranes were washed three times with TBS-T, before being incubated 1h at room temperature with HRP-conjugated secondary antibodies (1:2 000 in TBS-T). After three washing steps, ECL substrate was applied and HRP-mediated signals were visualized with the ChemiDoc Touch Imaging System (Bio-Rad).

1. Maier, J., B. Traenkle, and U. Rothbauer, *Real-time analysis of epithelial-mesenchymal transition using fluorescent single-domain antibodies.* Scientific Reports, 2015. **5**(1): p. 13402.
